# pre-miRBench: a multispecies benchmark and reference model for animal precursor microRNA prediction

**DOI:** 10.64898/2026.09.30.755660

**Authors:** Dimosthenis Tzimotoudis, Vasileios K. Chatzitolios, Panagiotis Alexiou, Georgios K. Georgakilas

## Abstract

**Motivation:** Precursor microRNA (pre-miRNA) prediction is used to prioritize genomic hairpins for miRNA annotation. Published predictors are trained and evaluated on different species collections, negative sets and data partitions, making reported gains difficult to compare.

**Results:** We introduce pre-miRBench, a benchmark of 7,056 200-nt windows centered on curated pre-miRNA loci and 70,560 genomic negatives with predicted hairpin structures from 71 animal species, with a fixed 1:10 positive-to-negative ratio in every partition. Four test sets cross whether the species and miRNA family are represented in training: known species/known family, known species/held-out family, held-out species/known family, and held-out species/held-out family. We retrained six published pre-miRNA predictors (DeepMir, deepMiRGene, dnnPreMiR, mirDNN, miRe2e and MuStARD) under the same benchmark splits and evaluated them alongside the pre-miRBench model, an ensemble of three neural networks developed with Agentomics, an autonomous LLM-agent system for biomedical machine learning. The pre-miRBench model achieved the highest mean average precision (AP) across the four test sets (0.9809; dnnPreMiR, 0.9719) and led three tests. The two held-out species were *Gallus gallus* and *Drosophila melanogaster*, and the joint species-and-family holdout contained 69 positives and 690 negatives. dnnPreMiR had the highest AP on the joint species-and-family holdout (0.9677 versus 0.9659). The AP difference (pre-miRBench minus dnnPreMiR) was −0.0018, with a paired bootstrap 95% confidence interval of −0.0445 to 0.0341. pre-miRBench establishes a common evaluation for species and miRNA-family holdouts and provides a reproducible reference model for future comparisons.

**Availability and implementation:** Reproducible source code is available at https://github.com/dimostzim/pre-miRBench. Datasets, trained model weights, record-level predictions, evaluation metrics and species-panel metadata for pre-miRBench and the six retrained predictors are available at https://zenodo.org/records/22813044.

## 1. Introduction

MicroRNAs (miRNAs) are short regulatory RNAs produced from longer transcripts that form characteristic hairpin structures. The discovery of *lin-4, let-7* and other miRNAs established their role in post-transcriptional regulation [1–4]. In canonical biogenesis, primary transcripts are processed into precursor hairpins and then into mature miRNAs that guide Argonaute-containing complexes to target RNAs [5–8]. Although small-RNA sequencing provides direct evidence for expressed products, computational prediction of precursor loci remains useful when expression is restricted to particular tissues or conditions, when only genome assemblies are available, and when prioritizing candidate loci for experimental analysis. Precursor predictors can also help prioritize candidate miRNA loci among regions enriched in small-RNA sequencing reads [9]. Direct comparison of pre-miRNA predictors is difficult because studies differ in annotation sources, negative sets, sequence lengths, available features, species composition and train–test partitions.

The apparent simplicity of a pre-miRNA hairpin masks a difficult discrimination problem. Millions of genomic sequences can potentially fold into pre-miRNA-like stem-loop structures, yet only a small subset are actually processed into miRNAs. Early prediction systems combined sequence composition, thermodynamic stability and structural descriptors using support vector machines or random forests [10– More recent methods learn representations from sequence or predicted structure with neural networks [9, 15–19].

Previous benchmarks have examined different aspects of pre-miRNA prediction. Stegmayer et al. compared learning approaches on human and Arabidopsis datasets derived from miRBase v17 while varying class ratios [20]. Bugnon et al. used miRBase v21 to label genome-derived hairpins from five species, treated the remaining hairpins as unlabeled, and evaluated each species separately by cross-validation [21]. Lopes et al. tested transfer across 45 species using miRBase v20 positives and pseudo-hairpins from coding sequences or pseudogenes, with within-species redundancy removed by clustering at 80% sequence similarity [22]. These studies reported public datasets or reproduction materials, but their described partitions did not cross species holdouts with named miRNA-family holdouts [20–22]. pre-miRBench combines that split design with curated MirGeneDB annotations, genomic negatives selected after hairpin filtering and classifier-based mining, and exact duplicate removal at the shortest model-input length.

Train–test partitions determine what type of generalization is evaluated. A random record-level split measures performance on new records from the same mixture of species and miRNA families used for training. Species holdouts test predictions for species absent from training, whereas family holdouts test predictions for miRNA families absent from training. Because homologous families occur across taxa, holding out a species does not necessarily remove related precursors from the training data [23–25]. Placing identical inputs in training and test can inflate performance estimates because the test example has already been seen during training. Biased resampling and incomplete reporting can also produce optimistic or difficult-to-reproduce estimates [26–28].

Negative-set construction and class prevalence also affect reported performance. Random genomic fragments often lack precursor-like hairpins, making them easy to distinguish from genuine pre-miRNAs. Small, hand-selected pseudo-hairpin sets may instead represent only a narrow subset of the candidates encountered during genome scanning [20, 21, 29]. Because true precursor loci are rare, accuracy can remain high even when a model produces many false-positive predictions. Precision– recall metrics, including average precision, are more informative under this imbalance because they reflect the trade-off between recovering positives and limiting false-positive predictions [30– 32]. Reviews and comparative studies have repeatedly identified negative-set construction, class balance and cross-species evaluation as barriers to direct comparison of pre-miRNA predictors [20–22, 33].

Reliable annotation is equally important. miRBase provides broad coverage and remains a major community resource [34], but historical miRNA collections vary in evidential support and annotation practice. Uniform annotation criteria were proposed early in the field [35], and MirGeneDB was developed to emphasize curated, evolutionarily supported miRNA genes with consistent family nomenclature [23, 24]. Its expanded taxonomic sampling provides an opportunity to evaluate generalization across both evolutionary lineages and miRNA families [25].

Here, we introduce pre-miRBench, a multispecies benchmark and model for pre-miRNA prediction. The task is to distinguish 200-nt windows centered on annotated pre-miRNA loci from other genomic windows with predicted hairpin structures. Locating new loci across a genome and experimentally confirming miRNA processing are outside the scope of this benchmark. We construct a 71-species dataset from curated MirGeneDB precursor loci and hard genomic negatives that already satisfy basic hairpin criteria. Four test sets distinguish predictions for species and miRNA families represented during training from predictions for held-out species and families. We retrain six published neural predictors under the same data partitions, class ratio and validation policy and evaluate them alongside a pre-miRNA prediction model developed with Agentomics [36]. We ask how predictor rankings change across the four test settings and how the pre-miRBench model compares with published predictors under the same benchmark partitions and evaluation protocol.

## 2. Materials and methods

### 2.1. Species panel, genome resources and precursor annotations

The benchmark included 71 of the 114 species in MirGeneDB 3.0 for which we could identify a reference genome assembly that matched the annotated precursor sequences and coordinates. Genome assemblies were obtained from NCBI Datasets (46 species), UCSC (22), Ensembl or Ensembl Metazoa (2), and the NHGRI Hydra portal (1). Assembly coordinates and chromosome names were checked against the corresponding source records [37–39]. Assembly identifiers, source URLs and SHA-256 checksums for the genome sequences and MirGeneDB annotation files are provided in the input manifest archived on Zenodo.

MirGeneDB browser export files were filtered to precursor entries. The raw manifest contained 12,053 precursor BED rows, all of which matched an assembly coordinate. Genomic intervals were manipulated using BEDTools-compatible operations [40]. For each locus, a strand-correct 200-nt window centered on the annotated precursor was extracted and converted to the RNA alphabet. Windows extending beyond a chromosome boundary were excluded. We excluded windows when lowercase soft-masked bases and uppercase N characters together accounted for more than 10% of the 200-nt sequence. Secondary structures and minimum free energies (MFEs) were computed with RNAfold 2.7.2 [41].

Of the 12,053 coordinate-matched rows, 14 were removed at chromosome boundaries and 3,724 (30.9%) exceeded the masked-base threshold, leaving 8,315 positive records after sequence-level quality filters. Exact duplicate handling was then applied at the model-input level as described below. A total of 1,259 redundant positive inputs were excluded, producing 7,056 final positive examples. Species-level counts for the final positive windows are included in the Zenodo archive.

Family identifiers were derived from MirGeneDB gene names. Species prefixes and locus or paralogue suffixes were normalized and miRNA names were mapped to their family stem. The resulting assignments retain the resource nomenclature and serve only to construct benchmark partitions.

### 2.2. Genomic candidate generation and hard-negative mining

Negative candidates were sampled from the same assemblies as the positive loci. Each genome was scanned on both strands with 200-nt windows at a 50-nt stride. Windows overlapping an annotated pre-miRNA interval were discarded. Windows with more than 10% lowercase soft-masked bases and uppercase N characters were removed. The scan was deterministic: the 50,000-window target was divided equally among chromosomes containing retained positive loci, and eligible windows were retained in coordinate order until each chromosome’s rounded-up quota was reached. Overlap operations used BEDTools-compatible interval logic [40].

Candidate windows were folded with RNAfold and retained only if they passed four deliberately permissive hairpin filters: MFE at most *−*10 kcal/mol, paired-nucleotide fraction of at least 0.40, a maximum contiguous stem of at least 8 base pairs, and a maximum unpaired run of at most 25 nt. These filters remove obviously non-hairpin sequences without defining a high-confidence miRNA class. Across the panel, 3,142,826 candidate windows were folded and 2,752,350 passed the hairpin filters. During genomic scanning, 6,165 overlapping windows and 9,588,347 repeat-rich or ambiguous windows were skipped. To reduce the abundance of easily rejected negatives, we performed four rounds of per-species hard-negative mining. Each candidate was represented by MFE, the frequencies of the 16 dinucleotides, maximum stem length, maximum loop length, bulge count, paired fraction and base-composition entropy. For each of the 69 represented species, a 10-member random-forest ensemble with 200 trees per member and class-balanced fitting was trained using only that species’ training positives [42]. For the two held-out species, the ensemble was trained using the pooled positives from the training partition; no validation or test positives were used for mining. At each round, a candidate was designated hard if at least half of the ensemble members assigned a positive-class probability of 0.5 or greater. Candidates were ordered first by the round in which they were identified and then by decreasing ensemble score; any remaining quota was filled from the highest-scoring unused pool. Random states were derived deterministically from seed 42 and the species index.

Final negative sampling was performed within species and split to preserve a 1:10 positive-to-negative ratio. The fixed ratio holds class prevalence constant across test settings. The selected negatives resemble precursor hairpins according to the stated folding criteria, lack overlap with the curated precursor set and were selected to challenge a sequence/structure classifier.

### 2.3. Split design and duplicate removal

The split design crosses two factors: whether a species is represented in training and whether a miRNA family is represented in training (Figure 1). *Gallus gallus* and *Drosophila melanogaster* were held out as widely used vertebrate and invertebrate model species. The remaining 69 species contributed to training and validation and to the known-species test partitions. Within the known-species panel, selected family groups were reserved completely from training to form the held-out-family test set. The fourth test set contains records from the two held-out species whose families were also absent from training.

**Figure 1.**
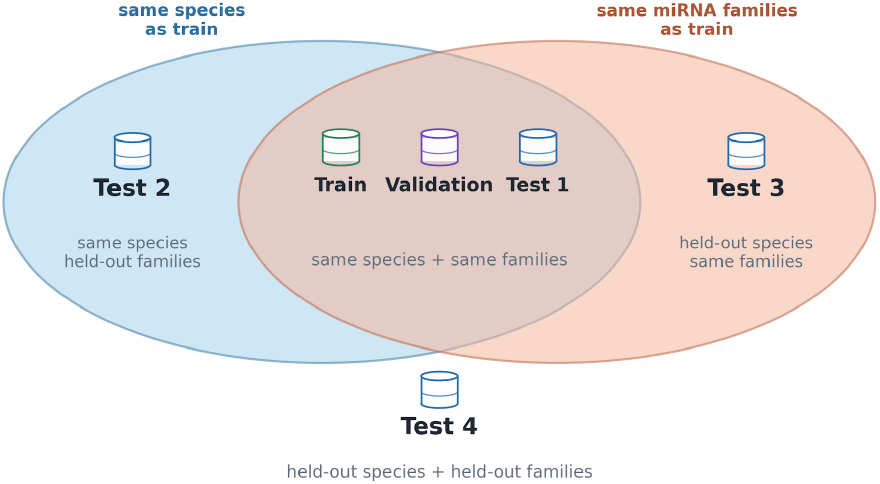
pre-miRBench split design. Training and validation contain species and miRNA families represented during training. The four test sets cross species status (represented or held out) with miRNA-family status (represented or held out)s.

Known-family validation and test examples were assigned as precursor-sequence groups, with at least one group from each represented family retained in training. Held-out families were assigned in their entirety. Using seed 42, precursor groups were sampled until validation and the known-species/known-family test each reached a target of 10% of eligible known-species positive records. Entire families were sampled for the known-species / held-out-family test until its target of 10% of eligible known-species positive records was reached. Because groups were kept intact and duplicates were removed later, final counts can differ from these targets. Validation contained only families represented in training.

Models in the comparison accept inputs ranging from 100 to 200 nt. To identify repeated model inputs consistently, we normalized and center-cropped every record to the shortest input length of 100 nt. Duplicate detection used this 100-nt key; models received their specified 100-, 160-, 180- or 200-nt inputs. Duplicate sequences were resolved after the biological split assignment, retaining records in the following order: the held-out-species / held-out-family test set, the other test sets, validation and training. The priority rule separates every test sequence from training and validation and can affect which records remain in a test set when duplicates exist. The benchmark contains 77,616 distinct 100-nt sequences, with no exact duplicate within or across labels or partitions.

The final training set contains 4,765 positives and 47,650 negatives from 69 species and 687 families (Table 1). Validation contains 631 positives and 6,310 negatives. The four test sets together contain 1,660 positives and 16,600 negatives. All sets preserve the 1:10 class ratio.

**Table 1.** Composition of the pre-miRBench partitions. Family counts refer to families represented among positive records.

| Partition | Species | Families | Positives | Negatives | Total |
| --- | --- | --- | --- | --- | --- |
| Train | 69 | 687 | 4,765 | 47,650 | 52,415 |
| Validation | 61 | 171 | 631 | 6,310 | 6,941 |
| Test 1 | 62 | 177 | 707 | 7,070 | 7,777 |
| Test 2 | 61 | 59 | 677 | 6,770 | 7,447 |
| Test 3 | 2 | 112 | 207 | 2,070 | 2,277 |
| Test 4 | 2 | 59 | 69 | 690 | 759 |
| Total | 71 | 790 | 7,056 | 70,560 | 77,616 |

Species/family status: Train, Validation and Test 1, known/known; Test 2, known/held out; Test 3, held out/known; Test 4, held out/held out.

### 2.4. Retraining published predictors

We retrained six published neural predictors with publicly available implementations or sufficiently specified model code: DeepMir [17], deepMiRGene [15], dnnPreMiR [18], mirDNN [16], miRe2e [19] and MuStARD [9]. The selected methods span sequence-only convolutional architectures, joint sequence/structure representations and multi-branch models.

Each tool was wrapped in a separate container and trained on the same pre-miRBench record identifiers. To preserve the published architecture, each adapter generated the model’s native centered crop from the common 200-nt window: 200 nt for DeepMir, 200 nt for deepMiRGene, 180 nt for dnnPreMiR, 160 nt for mirDNN, 100 nt for miRe2e and 200 nt for MuStARD. For positive samples, cropping retained the annotated precursor interval whenever its length permitted; negative samples were center-cropped. The 100-nt sequence served exclusively as the duplicate-detection key.

Model-specific feature generation followed the corresponding implementation. RNAfold-derived structure and MFE features were generated where required. MuStARD was run with its sequence and RNAfold branches; cross-species conservation tracks were omitted because they were unavailable uniformly across the 71 species [9]. miRe2e used its human-pretrained initialization for retraining, followed by inference with the newly trained structure, MFE and predictor components [19]. The shared protocol standardizes data and split exposure while preserving each method’s architecture, initialization and preprocessing.

The wrappers used validation AP for model selection where supported, with a 600-epoch cap and a patience of 20 for the epoch-based implementations. Tool-specific training loops were otherwise retained. Native batch sizes and architectural defaults were used: 128 for DeepMir, 256 for deepMiRGene, 200 for dnnPreMiR, 1,024 for mirDNN, and the implementation defaults for miRe2e and MuStARD. The four test sets were used only for final evaluation. Separate container definitions document the software and preprocessing setup used for each predictor.

### 2.5. pre-miRBench model

In addition to retraining the six published predictors, we developed the pre-miRBench model with Agentomics 1.0.4, an autonomous LLM-agent system for biomedical machine-learning experimentation [36], using gpt-5.6-sol through the OpenAI Codex provider. Agentomics operated on the fixed training and validation partitions, including 200-nt RNA sequences, RNAfold structures, MFEs and species codes, and produced the training and inference artifacts for the final model. Test labels were unavailable during development, candidate ensembles were limited to three components, and model selection used validation AP. Within this fixed task configuration, Agentomics performed model design, training, validation-based selection and ensemble construction without manual intervention. The complete run provenance log is released along with the produced model artifacts.

The selected predictor averages probabilities from one PairMessageCNN checkpoint and two SpeciesGraphGRU checkpoints. All three networks encode each 200-nt record with 12 position-wise channels representing the nucleotide, dot-bracket state, paired nucleotide and relative position of the paired base. PairMessageCNN uses a 64-channel convolutional stem followed by three residual blocks that combine local convolution with messages from paired positions at dilations 1, 2 and 4. Average and maximum pooled features are combined with standardized MFE before classification. SpeciesGraphGRU uses a 96-channel convolutional stem, three local and paired-position graph blocks, and a bidirectional GRU with 64 hidden units in each direction. Its classifier combines average, maximum and attention-pooled sequence features with a 16-dimensional species embedding and 13 training-standardized sequence and structure summaries. These summaries comprise MFE, GC and paired fractions, mean and maximum pair span, maximum nesting depth, maximum unpaired run and the fractions of six canonical and wobble pair orientations.

The PairMessageCNN and one SpeciesGraphGRU were trained with a reverse-complement view for half of the training records and were evaluated by averaging predictions for the original and reverse-complement views. The second SpeciesGraphGRU used the original view. During training of the species-conditioned models, 15% of species codes were replaced by an unknown-species index. All checkpoints used class-weighted binary cross-entropy, AdamW optimization, an exponential moving average of model parameters and early stopping by validation AP over at most 35 epochs. PairMessageCNN used a learning rate of 0.001 and batch size 512; SpeciesGraphGRU used a learning rate of 0.0005 and batch size 256. The final probability is the equal-weight mean of the three checkpoint probabilities, and binary predictions use a threshold of 0.5. The deployed ensemble contains 1,178,309 trainable parameters across its three checkpoints.

### 2.6. Evaluation

The seven predictors produced a score for every record in the four test sets, yielding 127,820 record-level predictions (18,260 records *×* seven methods) with no missing values. The primary metric was average precision (AP), computed from the complete ranked prediction list. AP summarizes precision at successive recall levels and differs from trapezoidal integration of the precision–recall curve. It is appropriate for the fixed 1:10 class ratio and emphasizes performance on the positive class [30, 31]. Secondary ranking performance was measured with area under the receiver-operating-characteristic curve (AUROC). At the fixed probability threshold of 0.5, we computed accuracy, precision, recall, specificity, F1 score and Matthews correlation coefficient (MCC). MCC was included because it summarizes all four cells of the binary confusion matrix and remains informative under unequal class frequencies [43].

Precision–recall curves were calculated separately for each predictor and test set. We summarized precision at the highest score threshold achieving at least 80% or 90% recall, and the maximum recall observed at precision of at least 90% or 95%. These points describe trade-offs within the test sets and were not used to select thresholds for future predictions.

Percentile 95% confidence intervals for AP and paired AP differences were estimated from 10,000 class-stratified bootstrap resamples of each test set (seed 42). Positive and negative records were resampled separately with replacement, using the same resampled indices for all models.

## 3. Results

### 3.1. Benchmarking and retraining models

The final benchmark contained 77,616 records from 71 species: 7,056 positives and 70,560 negatives. The final positive and negative records showed substantial overlap in the four RNAfold feature distributions (Figure 2).

**Figure 2.**
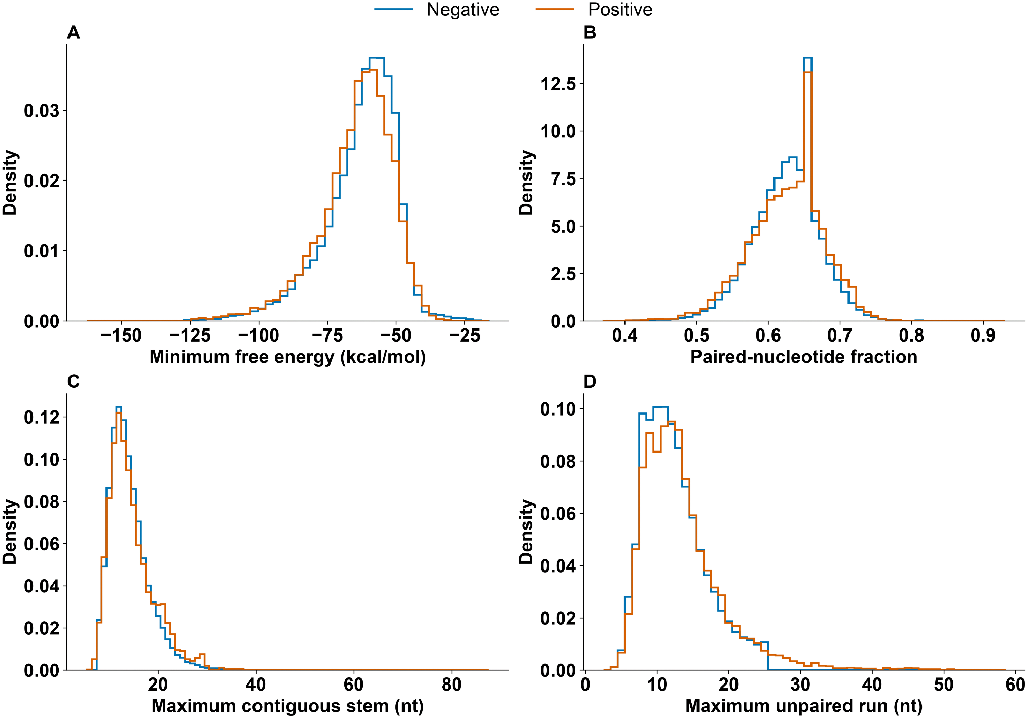
RNAfold feature distributions in the final benchmark. Distributions are shown separately for 7,056 positive and 70,560 negative records: (A) minimum free energy, (B) paired-nucleotide fraction, (C) maximum contiguous stem length and (D) maximum unpaired run.

Retraining the six published predictors on the fixed pre-miRBench splits produced different AP rankings across the four test settings (Table 2; Figure 3A). The pre-miRBench model ranked first on known species and known families (AP 0.9869), known species with held-out families (0.9813), and held-out species with known families (0.9895). dnnPreMiR ranked first on the joint species-and-family holdout (0.9677), followed by the pre-miRBench model (0.9659) and MuStARD (0.9526).

**Table 2.** Average precision across the four pre-miRBench test settings. Values are AP (95% bootstrap confidence interval). The highest AP in each column is bold.

| Predictor | Test 1 | Test 2 | Test 3 | Test 4 |
| --- | --- | --- | --- | --- |
| DeepMir | 0.8648 (0.8435–0.8858) | 0.8513 (0.8296–0.8722) | 0.8685 (0.8322–0.9031) | 0.8020 (0.7158–0.8833) |
| deepMiRGene | 0.9648 (0.9524–0.9760) | 0.9703 (0.9599–0.9800) | 0.9729 (0.9545–0.9881) | 0.9422 (0.8932–0.9820) |
| dnnPreMiR | 0.9782 (0.9699–0.9858) | 0.9595 (0.9478–0.9702) | 0.9821 (0.9678–0.9935) | **0.9677** (0.9415–0.9890) |
| mirDNN | 0.9845 (0.9772–0.9908) | 0.9561 (0.9439–0.9676) | 0.9853 (0.9738–0.9946) | 0.9113 (0.8548–0.9602) |
| miRe2e | 0.9628 (0.9511–0.9735) | 0.9499 (0.9365–0.9627) | 0.9532 (0.9303–0.9730) | 0.8685 (0.8027–0.9270) |
| MuStARD | 0.9815 (0.9745–0.9878) | 0.9696 (0.9595–0.9791) | 0.9789 (0.9632–0.9916) | 0.9526 (0.9137–0.9833) |
| pre-miRBench model | **0.9869** (0.9807–0.9925) | **0.9813** (0.9739–0.9880) | **0.9895** (0.9801–0.9966) | 0.9659 (0.9264–0.9955) |

Test 1: known species/known family; Test 2: known species/held-out family; Test 3: held-out species/known family; Test 4: held-out species/held-out family.

**Figure 3.**
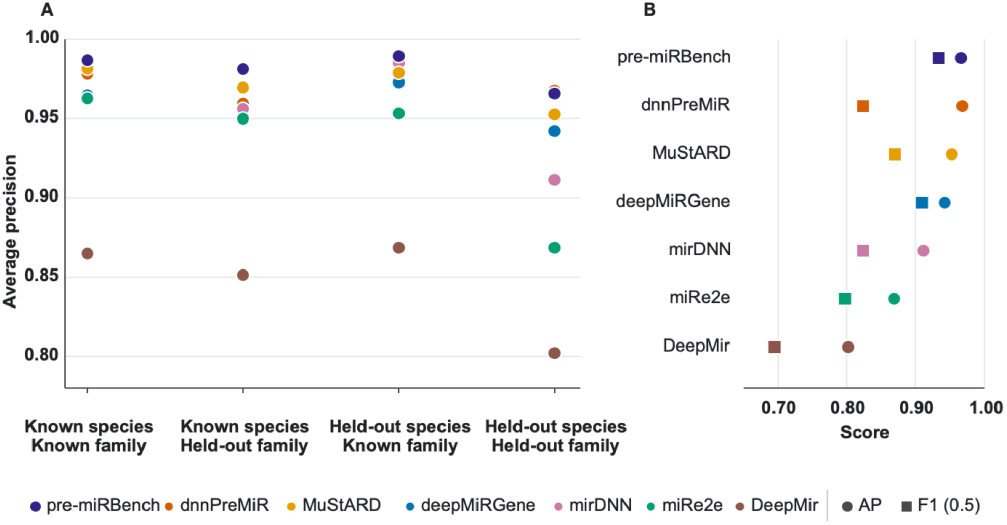
Predictor performance across four test settings. (A) Average precision for six retrained published methods and the pre-miRBench model on known-species/known-family, known-species/held-out-family, held-out-species/known-family and held-out-species/held-out-family test sets. (B) Average precision and F1 score at the fixed threshold of 0.5 on the held-out-species/held-out-family test.

Across the four tests, the pre-miRBench model had the highest mean AP (0.9809), followed by dnnPreMiR (0.9719) and MuStARD (0.9707). Holding out a species or family did not reduce AP uniformly: several predictors retained or improved AP on a single-axis holdout. The joint holdout produced the clearest change in ordering relative to the known-species/known-family test. On the held-out-species/held-out-family test, dnnPreMiR achieved AP 0.9677, the pre-miRBench model achieved 0.9659 and MuStARD achieved 0.9526. The AP difference between the pre-miRBench model and dnnPreMiR was *−*0.0018, with a paired bootstrap 95% confidence interval from *−*0.0445 to 0.0341.

The joint holdout contains 69 positives and 690 negatives from G. gallus and D. melanogaster. AP was lower in D. melanogaster for six of the seven predictors, including pre-miRBench (0.946 versus 0.975), dnnPreMiR (0.943 versus 0.980) and MuStARD (0.940 versus 0.957); DeepMir was similar between species (0.815 versus 0.811). These two species show within-test heterogeneity.

AP evaluates the full score ranking. Binary classification additionally requires a decision threshold. At the fixed threshold of 0.5, the pre-miRBench model achieved the highest accuracy (0.988), recall (0.913), F1 score (0.933) and MCC (0.927) on the joint holdout (Figure 3B; Table 3). dnnPreMiR, the AP leader on this test, achieved precision 0.980 and recall 0.710, with F1 0.824 and MCC 0.821.

**Table 3.** Performance on the held-out-species/held-out-family test at the fixed threshold of 0.5. AP is threshold-free and is repeated to show how the ranking differs from the binary classification metrics.

| Predictor | AP | Accuracy | Precision | Recall | Specificity | F1 | MCC |
| --- | --- | --- | --- | --- | --- | --- | --- |
| DeepMir | 0.8020 | 0.9526 | 0.8367 | 0.5942 | 0.9884 | 0.6949 | 0.6816 |
| deepMiRGene | 0.9422 | 0.9842 | 0.9524 | 0.8696 | 0.9957 | 0.9091 | 0.9016 |
| dnnPreMiR | 0.9677 | 0.9723 | 0.9800 | 0.7101 | 0.9986 | 0.8235 | 0.8213 |
| mirDNN | 0.9113 | 0.9723 | 0.9800 | 0.7101 | 0.9986 | 0.8235 | 0.8213 |
| miRe2e | 0.8685 | 0.9605 | 0.7468 | 0.8551 | 0.9710 | 0.7973 | 0.7777 |
| MuStARD | 0.9526 | 0.9763 | 0.8696 | 0.8696 | 0.9870 | 0.8696 | 0.8565 |
| pre-miRBench model | 0.9659 | 0.9881 | 0.9545 | 0.9130 | 0.9957 | 0.9333 | 0.9271 |

The equal 0.5-threshold confusion metrics of dnnPreMiR and mirDNN corresponded to AP values that differed by 0.056 on the same test. The pre-miRBench model recovered 63 of 69 positives with three false positives; dnnPreMiR recovered 49 positives with one false positive. Ranking quality and fixed-threshold classification therefore produced distinct model orderings. Precision–recall curves show how predictor performance changes across score thresholds in each test setting (Figure 4). At the highest recall attained while maintaining at least 95% precision on Test 4, pre-miRBench recovered 63 of 69 positives with three false positives (precision 0.955, recall 0.913), whereas dnnPreMiR recovered 53 positives with two false positives (precision 0.964, recall 0.768).

**Figure 4.**
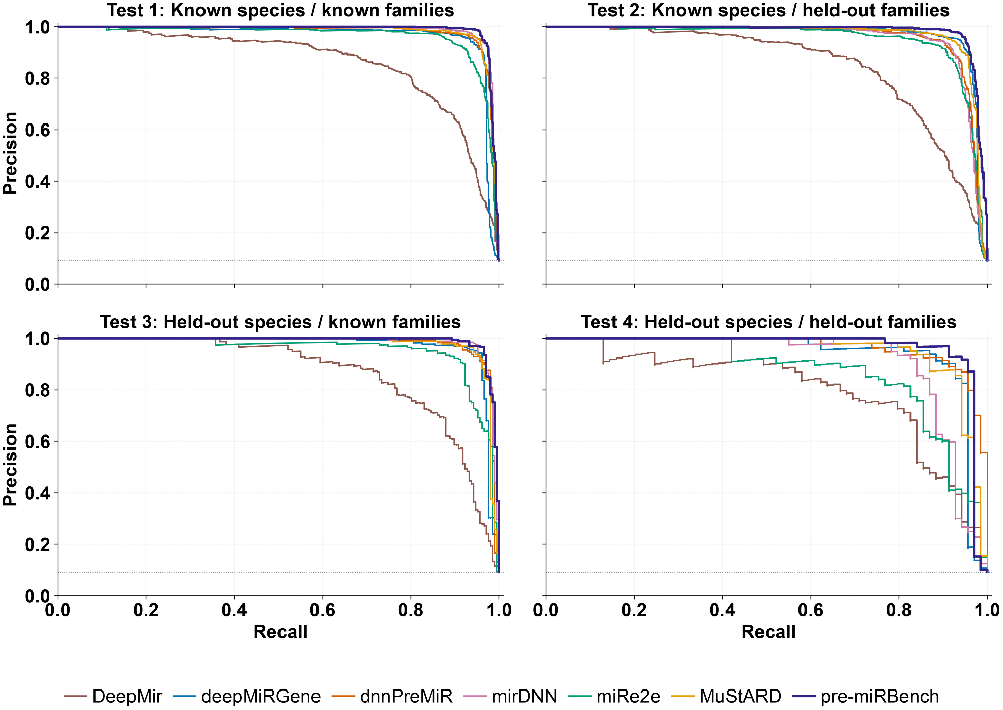
Precision–recall curves across the four test settings. Curves are shown for all seven predictors. The horizontal dotted line shows the positive prevalence (1/11).

## 4. Discussion

### 4.1. A shared benchmark changes model rankings

pre-miRBench places seven predictors on the same curated positives, precursor-like genomic negatives, class ratio and species-and-family partitions. The source methods were developed with different annotation collections, negative examples and split protocols [9, 15–19], so differences among their published scores can reflect dataset construction as well as model design. The common protocol retains each model’s input length and preprocessing while fixing which records are used for training, validation and testing. Hard-negative mining further changes the comparison by selecting genomic windows that already satisfy permissive hairpin criteria. Models are consequently evaluated against plausible genomic hairpins, which are more demanding controls than arbitrary sequence [20, 21, 29].

The shared evaluation changed the ordering of the six published predictors across the four tests. No published method led across settings. The pre-miRBench model achieved the highest mean AP and ranked first in three tests, while dnnPreMiR had the highest point estimate on the joint holdout. The AP difference (pre-miRBench minus dnnPreMiR) was *−*0.0018, with a paired bootstrap 95% confidence interval of *−*0.0445 to 0.0341. The joint-holdout data therefore support comparable ranking performance for the two leading models.

### 4.2. Model behavior differs across species and family holdouts

Species and family holdouts remove different sources of information from training. A species holdout can still contain miRNA families represented in other taxa, while a family holdout can contain species that contributed other precursors to training [23–25]. AP did not fall consistently when either factor was held out alone, and several predictors improved on one of the single-axis tests. These results indicate that absence from training along one biological axis does not by itself determine test difficulty. The retained family or species context and the composition of each test set also contribute.

All seven models had lower AP on the joint holdout than on the known-species/known-family test, but the size of the decrease differed markedly. Relative to the known-species/known-family test, AP decreased by 0.0105 for dnnPreMiR and 0.0210 for the pre-miRBench model, compared with 0.0732 for mirDNN and 0.0943 for miRe2e. The two held-out species also differed, as six of seven predictors had lower AP in *D. melanogaster* than in *G. gallus*. The joint holdout therefore reveals method-specific sensitivity to the combined change in species and family composition, which would not be visible from a single aggregate score.

### 4.3. Agentic model development produced a competitive reference model

Agentomics [36] developed the pre-miRBench model using the fixed training and validation partitions while test labels remained unavailable. The selected ensemble combines one PairMessageCNN and two SpeciesGraphGRU checkpoints. Their representations cover local sequence and structure patterns, interactions between paired positions, recurrent context across the complete window and global folding summaries. Two components also use species conditioning, with an unknown-species index presented during training, so the deployed model can process species absent from the training panel.

The Agentomics-derived model ranked within the top two in every test, providing direct evidence that the agentic workflow produced a competitive benchmark-native predictor under the fixed protocol. Its development also yielded complete training and inference programs, fitted preprocessing metadata, checkpoints and record-level predictions. Releasing these artifacts makes the selected model an inspectable reference and allows future predictors to be evaluated on the same records and evaluation code.

### 4.4. Benchmark scope and limitations

The computational negative class defines the scope of the reported scores. Selected windows lack overlap with MirGeneDB precursors and passed permissive hairpin filters, yet some may be unannotated miRNAs, other structured RNAs or condition-specific hairpins because absence from the annotation set does not establish absent miRNA processing. Performance therefore measures discrimination between curated precursors and these genomic controls. The false-discovery rate of whole-genome annotation remains unmeasured [20, 21, 29]. The fixed 1:10 ratio supports comparison across models and tests, but genome scanning is substantially more imbalanced and may yield more false-positive calls [30, 31].

Duplicate removal excluded identical 100-nt model inputs across partitions, but homologous non-identical precursors and longer shared subsequences may remain. Family assignments follow normalized MirGeneDB names [23–25] instead of sequence-identity clusters. The joint holdout contains 69 positives and 690 negatives from only *G. gallus* and *D. melanogaster*, and the confidence interval for the two leading models was wide relative to their AP difference. More held-out species and similarity-clustered families are needed to resolve small model differences and measure transfer across broader evolutionary distances.

Hard-negative mining uses RNAfold-derived features and a random-forest ensemble, which may favor or disadvantage predictors according to their use of related thermodynamic descriptors. The pre-miRBench model was selected on one validation partition, without component ablations or repeated Agentomics runs, leaving the contribution of each ensemble component and the stability of the selected design unresolved. The three checkpoints and reverse-complement averaging for two checkpoints also increase inference cost. Updated MirGeneDB annotations, experimental negative evidence, externally submitted predictions, component ablations and repeated development runs would extend the benchmark while retaining the fixed comparison established here.

## 5. Conclusions

pre-miRBench provides a 71-species dataset, four fixed species-and-family test settings, reproducible implementations for six published predictors and a new reference model. Under shared retraining, the pre-miRBench model achieved the highest mean AP and led three tests, while dnnPreMiR was marginally higher on the joint holdout. The fixed test sets, trained weights and record-level predictions provide a reproducible reference for evaluating future precursor-miRNA predictors.

## Data availability

The source code is available at https://github.com/dimostzim/pre-miRBench. For the pre-miRBench model, it also provides the deployment manifest, fitted preprocessing metadata, locked uv environment, and training, inference and evaluation programs. Datasets, trained model weights, record-level predictions and evaluation metrics for pre-miRBench and the six retrained predictors are available at https://zenodo.org/records/22813044.

## References

1. Lee RC, Feinbaum RL, Ambros V. The C. elegans heterochronic gene lin-4 encodes small RNAs with antisense complementarity to lin-14. Cell. 1993;75(5):843–854. doi:10.1016/0092-8674(93)90529-Y

2. Wightman B, Ha I, Ruvkun G. Posttranscriptional regulation of the heterochronic gene lin-14 by lin-4 mediates temporal pattern formation in C. elegans. Cell. 1993;75(5):855–862. doi:10.1016/0092-8674(93)90530-4

3. Reinhart BJ, Slack FJ, Basson M, Pasquinelli AE, Bettinger JC, Rougvie AE, et al. The 21-nucleotide let-7 RNA regulates developmental timing in Caenorhabditis elegans. Nature. 2000;403(6772):901–906. doi:10.1038/35002607

4. Lagos-Quintana M, Rauhut R, Lendeckel W, Tuschl T. Identification of novel genes coding for small expressed RNAs. Science. 2001;294(5543):853–858. doi:10.1126/science.1064921

5. Bartel DP. MicroRNAs: genomics, biogenesis, mechanism, and function. Cell. 2004;116(2):281–297. doi:10.1016/S0092-8674(04)00045-5

6. Ha M, Kim VN. Regulation of microRNA biogenesis. Nat Rev Mol Cell Biol. 2014;15(8):509–524. doi:10.1038/nrm3838

7. Bartel DP. Metazoan microRNAs. Cell. 2018;173(1):20–51. doi:10.1016/j.cell.2018.03.006

8. Treiber T, Treiber N, Meister G. Regulation of microRNA biogenesis and its crosstalk with other cellular pathways. Nat Rev Mol Cell Biol. 2019;20(1):5–20. doi:10.1038/s41580-018-0059-1

9. Georgakilas GK, Grioni A, Liakos KG, Chalupova E, Plessas FC, Alexiou P. Multi-branch convolutional neural network for identification of small non-coding RNA genomic loci. Sci Rep. 2020;10(1):9486. doi:10.1038/s41598-020-66454-3

10. Xue C, Li F, He T, Liu GP, Li Y, Zhang X. Classification of real and pseudo microRNA precursors using local structure-sequence features and support vector machine. BMC Bioinformatics. 2005;6(1):310. doi:10.1186/1471-2105-6-310

11. Jiang P, Wu H, Wang W, Ma W, Sun X, Lu Z. MiPred: classification of real and pseudo microRNA precursors using random forest prediction model with combined features. Nucleic Acids Res. 2007;35(Web Server issue):W339–W344. doi:10.1093/nar/gkm368

12. Yousef M, Jung S, Showe LC, Showe MK. Learning from positive examples when the negative class is undetermined— microRNA gene identification. Algorithms Mol Biol. 2008;3(1):2. doi:10.1186/1748-7188-3-2

13. Batuwita R, Palade V. microPred: effective classification of pre-miRNAs for human miRNA gene prediction. Bioinformatics. 2009;25(8):989–995. doi:10.1093/bioinformatics/btp107

14. Tran VDT, Tempel S, Zerath B, Zehraoui F, Tahi F. miRBoost: boosting support vector machines for microRNA precursor classification. RNA. 2015;21(5):775–785. doi:10.1261/rna.043612.113

15. Park S, Min S, Choi HS, Yoon S. Deep recurrent neural network-based identification of precursor microRNAs. In: Advances in Neural Information Processing Systems 30; 2017.

16. Yones C, Raad J, Bugnon LA, Milone DH, Stegmayer G. High precision in microRNA prediction: a novel genome-wide approach with convolutional deep residual networks. Comput Biol Med. 2021;134:104448. doi:10.1016/j.compbiomed.2021.104448

17. Cordero J, Menkovski V, Allmer J. Detection of pre-microRNA with convolutional neural networks. bioRxiv [Preprint]. 2019. doi:10.1101/840579

18. Zheng X, Fu X, Wang K, Wang M. Deep neural networks for human microRNA precursor detection. BMC Bioinformatics. 2020;21(1):17. doi:10.1186/s12859-020-3339-7

19. Raad J, Bugnon LA, Milone DH, Stegmayer G. miRe2e: a full end-to-end deep model based on transformers for prediction of pre-miRNAs. Bioinformatics. 2022;38(5):1191–1197. doi:10.1093/bioinformatics/btab823

20. Stegmayer G, Di Persia LE, Rubiolo M, Gerard M, Pividori M, Yones C, et al. Predicting novel microRNA: a comprehensive comparison of machine learning approaches. Brief Bioinform. 2019;20(5):1607–1620. doi:10.1093/bib/bby037

21. Bugnon LA, Yones C, Milone DH, Stegmayer G. Genome-wide discovery of pre-miRNAs: comparison of recent approaches based on machine learning. Brief Bioinform. 2021;22(3):bbaa184. doi:10.1093/bib/bbaa184

22. Lopes IDON, Schliep A, de Carvalho Apdlf. Automatic learning of pre-miRNAs from different species. BMC Bioinformatics. 2016;17(1):224. doi:10.1186/s12859-016-1036-3

23. Fromm B, Billipp T, Peck LE, Johansen M, Tarver JE, King BL, et al. A uniform system for the annotation of vertebrate microRNA genes and the evolution of the human microRNAome. Annu Rev Genet. 2015;49(1):213–242. doi:10.1146/annurev-genet-120213-092023

24. Fromm B, Høye E, Domanska D, Zhong X, Aparicio-Puerta E, Ovchinnikov V, et al. MirGeneDB 2.1: toward a complete sampling of all major animal phyla. Nucleic Acids Res. 2022;50(D1):D204– D210. doi:10.1093/nar/gkab1101

25. Clarke AW, Høye E, Hembrom AA, Paynter VM, Vinther J, Wyrożemski Ł, et al. MirGeneDB 3.0: improved taxonomic sampling, uniform nomenclature of novel conserved microRNA families and updated covariance models. Nucleic Acids Res. 2025;53(D1):D116–D128. doi:10.1093/nar/gkae1094

26. Varma S, Simon R. Bias in error estimation when using cross-validation for model selection. BMC Bioinformatics. 2006;7(1):91. doi:10.1186/1471-2105-7-91

27. Kapoor S, Narayanan A. Leakage and the reproducibility crisis in machine-learning-based science. Patterns (N Y). 2023;4(9):100804. doi:10.1016/j.patter.2023.100804

28. Walsh I, Fishman D, Garcia-Gasulla D, Titma T, Pollastri G, ELIXIR Machine Learning Focus Group, et al. DOME: recommendations for supervised machine learning validation in biology. Nat Methods. 2021;18(10):1122–1127. doi:10.1038/s41592-021-01205-4

29. Ritchie W, Gao D, Rasko JEJ. Defining and providing robust controls for microRNA prediction. Bioinformatics. 2012;28(8):1058–1061. doi:10.1093/bioinformatics/bts114

30. Davis J, Goadrich M. The relationship between precision-recall and ROC curves. In: Proceedings of the 23rd International Conference on Machine Learning; 2006. p. 233–240. doi:10.1145/1143844.1143874

31. Saito T, Rehmsmeier M. The precision-recall plot is more informative than the ROC plot when evaluating binary classifiers on imbalanced datasets. PLoS One. 2015;10(3):e0118432. doi:10.1371/journal.pone.0118432

32. Zhong Y, Xuan P, Han K, Zhang W, Li J. Improved pre-miRNA classification by reducing the effect of class imbalance. Biomed Res Int. 2015;2015:960108. doi:10.1155/2015/960108

33. Yones C, Stegmayer G, Milone DH. Genome-wide pre-miRNA discovery from few labeled examples. Bioinformatics. 2018;34(4):541–549. doi:10.1093/bioinformatics/btx612

34. Kozomara A, Birgaoanu M, Griffiths-Jones S. miRBase: from microRNA sequences to function. Nucleic Acids Res. 2019;47(D1):D155–D162. doi:10.1093/nar/gky1141

35. Ambros V, Bartel B, Bartel DP, Burge CB, Carrington JC, Chen X, et al. A uniform system for microRNA annotation. RNA. 2003;9(3):277–279. doi:10.1261/rna.2183803

36. Martinek V, Gariboldi A, Tzimotoudis D, Galea M, Zacharopoulou E, Escudero AA, et al. Agentomics: an agentic system that autonomously develops novel state-of-the-art solutions for biomedical machine learning tasks. Bioinformatics. 2026;42(Suppl 1):btag250. doi:10.1093/bioinformatics/btag250

37. O’Leary NA, Cox E, Holmes JB, Anderson WR, Falk R, Hem V, et al. Exploring and retrieving sequence and metadata for species across the tree of life with NCBI Datasets. Sci Data. 2024;11(1):732. doi:10.1038/s41597-024-03571-y

38. Cunningham F, Allen JE, Allen J, Alvarez-Jarreta J, Amode MR, Armean IM, et al. Ensembl 2022. Nucleic Acids Res. 2022;50(D1):D988–D995. doi:10.1093/nar/gkab1049

39. Kent WJ, Sugnet CW, Furey TS, Roskin KM, Pringle TH, Zahler AM, et al. The human genome browser at UCSC. Genome Res. 2002;12(6):996–1006. doi:10.1101/gr.229102

40. Quinlan AR, Hall IM. BEDTools: a flexible suite of utilities for comparing genomic features. Bioinformatics. 2010;26(6):841–842. doi:10.1093/bioinformatics/btq033

41. Lorenz R, Bernhart SH, Höner zu Siederdissen C, Tafer H, Flamm C, Stadler PF, et al. ViennaRNA Package 2.0. Algorithms Mol Biol. 2011;6(1):26. doi:10.1186/1748-7188-6-26

42. Breiman L. Random forests. Mach Learn. 2001;45(1):5–32. doi:10.1023/A:1010933404324

43. Chicco D, Jurman G. The advantages of the Matthews correlation coefficient (MCC) over F1 score and accuracy in binary classification evaluation. BMC Genomics. 2020;21(1):6. doi:10.1186/s12864-019-6413-7

